# Quantifying the geographic mosaic of coevolutionary temperature: from coldspots to hotspots

**DOI:** 10.1101/2025.11.27.691007

**Authors:** Klementyna A. Gawecka, Fernando Pedraza, Cecilia S. de Andreazzi, Jordi Bascompte

**Affiliations:** Department of Evolutionary Biology and Environmental Studies, University of Zurich, Winterthurerstrasse 190, 8057 Zürich, Switzerland; UK Centre for Ecology & Hydrology, Bush Estate, Penicuik, EH26 0QB, UK; Departamento de Biodiversidad, Ecología y Evolución, Universidad Complutense de Madrid, Av. Complutense, 28040 Madrid, Spain

## Abstract

The Geographic Mosaic Theory of Coevolution (GMTC) predicts that reciprocal evolutionary effects vary across landscapes, generating hotspots and coldspots. Traditionally, these states are treated as discrete categories, even though the intensity of coevolutionary selection can vary continuously. To capture this variation, we introduce a concept of coevolutionary temperature, ranging from coldspot to hotspot. We propose two complementary metrics to quantify it: reciprocity and strength of pairwise evolutionary effects. We also extend the GMTC framework beyond its traditional focus on pairwise systems to species-rich communities. Applying this approach to empirical plant-pollinator networks in a fragmented landscape, we find pronounced geographic mosaics in coevolutionary temperature. Smaller habitat patches support small, highly connected, and weakly nested communities with high reciprocity and strength, suggesting that they act as coevolutionary hotspots. In contrast, larger patches host species-rich, poorly connected, and highly nested communities with low reciprocity and strength, consistent with coldspots. At the interaction scale, reciprocity depends on degree similarity, with interactions between species that have similar numbers of partners exhibiting higher reciprocity. Together, these results highlight the strong dependence of coevolutionary effects on spatial variation in community structure and show how extending the geographic mosaic framework to species-rich communities can deepen our understanding of coevolution in complex systems.

## Introduction

Coevolution is not a uniform process across space. Instead, the evolutionary trajectories of interacting species can differ among locations, resulting in geographic variation in traits. For example, both the flowers of *Lithophragma* species and their pollinators, *Greya* moths, show spatial differences in morphology (Thompson et al., 2017). Such variation arises because both local abiotic (e.g., environmental conditions) and biotic factors (e.g., species composition) influence the intensity and direction of natural selection. The Geographic Mosaic Theory of Coevolution (GMTC, Thompson, 2005) provides a framework for understanding these patterns by highlighting three processes. First, *geographic selection mosaics* arise when natural selection differs among populations due to spatial variation in how the fitness of one species depends on the genotypes of its interacting partner. Second, *trait remixing* continuously changes the genetic structure through mutation, gene flow, drift, and colonization–extinction dynamics. Third, *coevolutionary hotspots and coldspots* emerge because the intensity of reciprocal selection between interacting species varies across environments, with hotspots showing reciprocal selection and coldspots showing non-reciprocal selection.

While GMTC traditionally refers to hotspots and coldspots as discrete states, the intensity of selection can vary, producing a gradient from cold to hot (Thompson, 1999). An interaction may show no reciprocal selection (i.e., no coevolution), reciprocal but weak selection, strong reciprocal selection, or partly-reciprocal (strong on one but weak on the other) selection. Therefore, quantifying coevolutionary temperature in terms of continuous measures of *reciprocity* and *strength* may provide a more informative representation of coevolutionary effects. Yet, such measures have not been formally defined. Moreover, GMTC has been developed primarily for pairwise interactions, even though real communities contain many interacting species. This raises a key unresolved question: how does coevolutionary temperature vary within species-rich communities?

Despite their explanatory potential, most empirical studies of GMTC have focused on simple communities involving two or three species (e.g., Rusman et al., 2025b,a; Hague et al., 2020; Thompson and Cunningham, 2002; Benkman, 1999; Laine, 2009, and references therein). Extending this framework to species-rich communities poses challenges (but see Erdos et al., 2025; Hall et al., 2020). Reciprocal evolutionary effects are difficult to quantify in complex communities, and disentangling the roles of abiotic versus biotic drivers of coevolution is rarely feasible (Yoder, 2025; Nuismer et al., 2022; terHorst et al., 2018). Theoretical models have helped address some of these challenges by integrating network and coevolutionary theory (Assis et al., 2020; Andreazzi et al., 2017; Guimarães et al., 2017; Nuismer et al., 2013). Within the GMTC context, such models have been applied to explore the consequences of spatially distributed hotspots and coldspots (Cosmo et al., 2024; Fernandes et al., 2019; Medeiros et al., 2018; Gibert et al., 2013; Gomulkiewicz et al., 2000). However, important questions about the emergence of hotspots and coldspots remain unresolved.

The GTMC is becoming increasingly relevant in this time of rapid global change, especially due to the widespread loss and fragmentation of natural habitats (Zou et al., 2025; Haddad et al., 2015). Fragmented landscapes form mosaics of patches that differ in area and connectivity, as well as in community composition and structure (Wang et al., 2023; Li et al., 2022; Librán-Embid et al., 2021; Jauker et al., 2019; Emer et al., 2018; Grass et al., 2018; Gilarranz et al., 2015). Such spatial heterogeneity can generate selection mosaics and influence trait remixing (e.g., Assis et al., 2022; Gowda and Kress, 2013; Gómez et al., 2009). Consequently, coevolutionary temperature may vary across patches within the same landscape. However, the drivers of this spatial variation remain unclear (Agrawal and Zhang, 2021). In particular, we do not yet know which habitat patches, and which pairwise interactions, are most likely to function as hotspots or coldspots.

Here, we propose quantifying *coevolutionary temperature* in terms of two complementary, continuous measures of pairwise coevolutionary effects: *reciprocity* and *strength* (Figure 1). We apply this approach to empirical plant–pollinator communities in fragmented grasslands (Jauker et al., 2019; Grass et al., 2018) and use a coevolutionary model (Guimarães et al., 2017) to quantify temperature at both community and interaction scales. We assess how patch area and local network structure affect reciprocity and strength. This community-level perspective allows us to determine not only which habitat patches are most likely to harbour hotspots, but also which interactions within a community are most likely to experience strong reciprocal coevolutionary effects.

**Figure 1:**
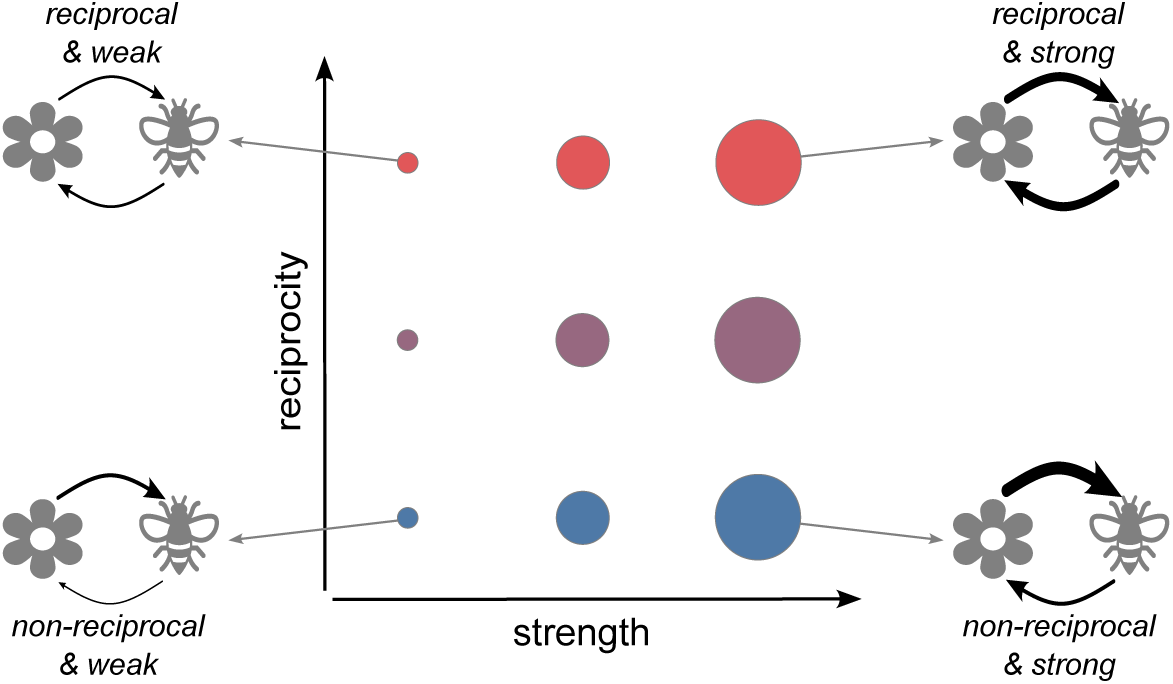
Our continuous measures of coevolutionary temperature: reciprocity and strength. Reciprocity (indicated by colour gradient from blue to red) compares the similarity of magnitudes of evolutionary effects of species *i* on species *j* and vice versa (thickness of arrows between plant and pollinator). Coevolution is highly reciprocal if the two effects are similar in magnitude. Strength (indicated by circle size) quantifies the average evolutionary effect for a pairwise interaction. Coevolution is strong if both effects are high.

## Methods

### Empirical dataset

As our case study, we focused on fragmented calcareous grasslands in Germany (Jauker et al., 2019; Grass et al., 2018). Out of the 285 grasslands fragments present in the region, 32 were chosen as study sites where plant-pollinator interactions were observed. These patches range in size from 314 to 51,395 m^2^. The observed bipartite plant-pollinator networks vary widely in size and structure (Figures S1-S2). For more information on the empirical dataset, refer to Grass et al. (2018) and Jauker et al. (2019).

### Coevolutionary model

We used the model proposed by Guimarães et al. (2017) to simulate coevolution between the species observed in each grassland patch. This model is based on a selection gradient which connects the evolution of a single trait with the mean fitness consequences of interactions between species and with the environment. The mean trait value of species *i* over a time step, *t*, is given by:

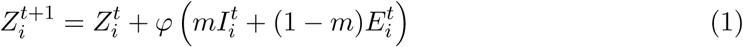

where *φ* is a compound parameter that affects the slope of the selection gradient and is proportional to the additive genetic variance, 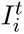 and 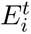 are the partial selection differentials corresponding to selection imposed by species interactions and environment, respectively, and *m* is the relative importance of interactions in trait evolution. Note that, when *m* = 0 or *m* = 1, trait evolution is entirely governed by the environment or species interactions, respectively.

The trait change driven by species interactions, 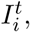 is defined as:

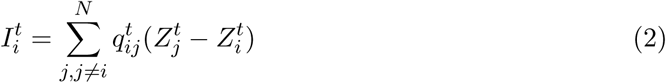

where *N* is the number of species in the local network, and 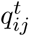 is an element of the Q-matrix which describes the evolutionary effect of species *j* on species *i*. It is defined as:

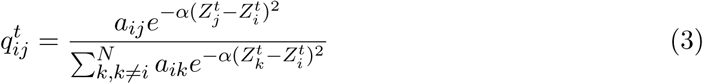

where *a_ij_* is an element of the binary adjacency matrix of the local interaction network (i.e., *a_ij_*= 1 indicates an interaction between species *i* and *j*), and *α* is a constant controlling the sensitivity to trait matching.

The trait change driven by the environment, 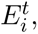 is defined as:

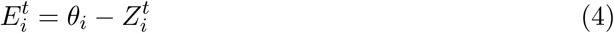

where *θ_i_* is the environmental optimum of species *i* (i.e. the phenotype favoured by the environmental selection).

In summary, the model tracks the evolution of a single trait for each species in the community. This trait evolves in response to both species interactions and environmental conditions. We assume that mutualistic interactions favour trait matching between partners. Regarding the environment, each species is assumed to have an optimal trait value toward which it evolves. Ultimately, trait evolution results from the combined influence of these two sources of selection.

We simulated the coevolutionary dynamics on each of the 32 empirical communities until a steady state, where species traits no longer change in time. For each community, we performed 100 replicate simulations which differed in species’ environmental optima (*θ_i_*). For each replicate, we sampled *θ_i_* from a uniform distribution between 0 and 10. We set the initial trait values to be equal to *θ_i_*, and *φ* to 0.5. However, note that while the initial trait values and *φ* influence the time to reach steady state, they have no effect on the steady state value of traits. We then averaged all quantities measured (see Coevolutionary temperature measure) across the 100 replicates. The results presented here were obtained with *α* = 0.1 and *m* = 0.5. Our findings are robust to *m* and *α* (see Figures S4-S6 where we varied *m* between 0.1 and 0.9, and *α* between 0.01 and 1).

### Coevolutionary temperature measures

We propose quantifying the coevolutionary temperature at the interaction scale in terms of two complementary continuous measures: *reciprocity* and *strength* of pairwise evolutionary effects (Figure 1). In our coevolutionary model, the pairwise evolutionary effects of species *j* on species *i*, *q_ij_*, are encapsulated by the Q-matrix (Equation 3). Thus, we compute both reciprocity and strength from the elements of the Q-matrix at the steady state.

We defined reciprocity, *R_ij_*, between species *i* and *j* as:

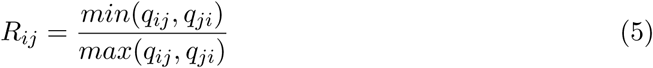

Dividing the smaller evolutionary effect by the larger one ensures that the measure of reciprocity is bound between 0 and 1. *R_ij_* = 0 indicates non-reciprocal selection where only one of the two species has an effect on the other. Conversely, *R_ij_* = 1, corresponds to a fully reciprocal selection where both species affect each other equally. Note that our measure of reciprocity is analogous to the asymmetry of pairwise interaction for weighted interaction networks proposed by Bascompte et al. (2006).

Strength, *S_ij_*, is defined as the average evolutionary effect that a pair of species *i* and *j* have on each other:

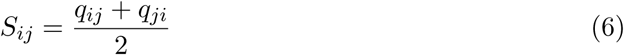

*S_ij_*is also bound between 0 and 1. Values close to 0 indicating weak (or non-existent) evolutionary effects, whereas values close to 1 correspond to strong evolutionary effects. Note that our definition of strength differs from that proposed by Bascompte et al. (2006) which is a quantitive extension of species degree.

We measured reciprocity and strength for each interaction in all local networks. Additionally, we studied the coevolutionary temperature at the community scale by averaging both reciprocity and strength across all interaction in each local network.

### Data analysis

We analysed the local network structure in terms of the number of species, number of interactions, connectance, nestedness and modularity. Connectance is the proportion of possible interactions that are realised in the local network. Nestedness is a pattern whereby a core group of generalist species interacts with other generalists and specialist, whereas specialists interact primarily with the generalists. We quantified nestedness using the measure proposed by Fortuna et al. (2019). Modularity measures the division of a network into modules with many interactions within these modules and few interactions between modules. We used the ‘igraph’ package in R with a multilevel modularity optimisation algorithm to quantify modularity (Blondel et al., 2008). As both nestedness and modularity are affected by network size, we standardised the computed values as Z-scores using a probabilistic cell null model (Bascompte et al., 2003).

To investigate the relationships between patch area, local network structure and co-evolutionary temperature measures, we used a Structural Equation Model (SEM). Based on previous analysis of the empirical dataset (Grass et al., 2018; Jauker et al., 2019), we expected patch size to affect the local network structure. We postulated that the local network structure, in turn, affects both the average reciprocity and the average strength across all interactions in the local network. Our analysis accounted for correlations between network structure measures in two steps. First, we summarised the number of species, number of interactions and connectance using a principal component analysis (PCA). The first principal component (PC1) explained 90% of variance and was positively correlated with the number of species (99%) and number of interactions (95%), and negatively correlated with connectance (-90%). Thus, higher PC1 values more species- and interaction-rich but less connected networks. Second, to account for residual correlations among network properties, our structural equation model included covariance terms between PC1, standardised nestedness and standardised modularity (double-headed arrows in Figure 3). In the SEM, we modelled the effects of patch area (*ln*-transformed) on local network structure (PC1, nestedness and modularity), and subsequently on coevolutionary temperature measures (reciprocity and strength averaged across all local interactions) using linear models (Table S1). We performed SEM analysis using ‘piecewiseSEM’ R package (Lefcheck, 2016), and evaluated model fit using Fisher’s C and Chi-Squared statistics, both of which indicated adequate fit (p-values*>*0.05).

At the interaction level, we defined *interaction degree symmetry* as:

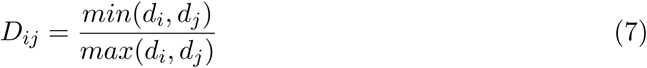

where *d_i_*is the number of interaction partners of species *i* (i.e., its degree). This measure ranges between 0 and 1, with values close to 0 corresponding to an interaction between species with very different degrees (i.e., a generalist and a specialist). At the other extreme, *D_ij_* = 1 indicates an interaction between species with the same number of partners. We related coevolutionary reciprocity and strength of interactions to their degree symmetry. To explore the relationship between the average interaction degree symmetry in a local network, local network structure and coevolutionary temperature, we constructed an additional SEM (see Figure S3, Table S2).

## Results

We find a geographic mosaic in coevolutionary temperature, defined in terms of reciprocity and strength, both at community and interaction scales. Spatial variation in local community size and structure across habitat fragments results in different average community reciprocity and strength (Figure 2A). Within local communities, there is also a large variation in coevolutionary temperature among pairwise interactions (Figure 2B). Moreover, the same pair of interacting species may have different reciprocity and strength in different fragments, depending on the local community. For example, the interaction between *Knautia arvensis* and *Episyrphus balteatus* (the most abundant interaction in this landscape) has relatively high reciprocity and strength in fragment 24, but low in fragment 15 (Figure 2B).

**Figure 2:**
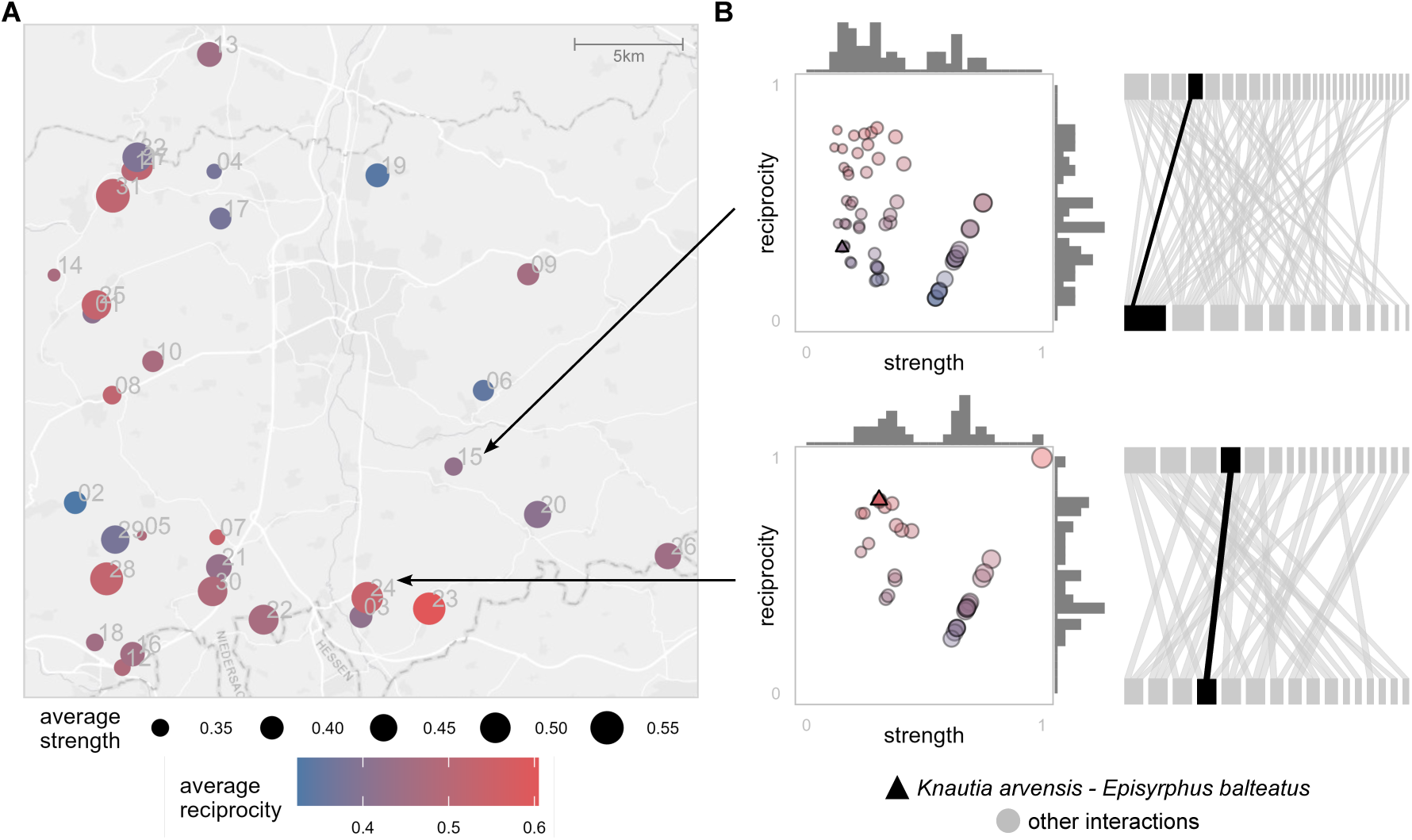
Geographic mosaic of coevolutionary temperature quantified in terms of reciprocity and strength at (A) community and (B) interaction scales. (A) The locations of the 32 calcareous grassland fragments studied. Point colour and size indicate reciprocity and strength, respectively, averaged across all observed local interactions. (B) The local communities in two selected grassland fragments. The left plots show reciprocity and strength of all local interactions. The right plots show the local interaction network between plants (bottom row) and pollinators (top row). The interaction between *Knautia arvensis* and *Episyrphus balteatus* (the most abundant interaction) is indicated by triangles and highlighted in black on the network, whereas all other interactions are shown as circles or in grey.

At the community scale, the average reciprocity and strength depend on the local network size and structure, which in turn is determined by habitat patch area (Figures 3, S2). Larger patches harbour more species, more interactions, less connected (i.e., high PC1 values) and more nested networks, than smaller patches. We find strong negative effects of: (1) nestedness on coevolutionary reciprocity and (2) PC1 on coevolutionary strength. This means that large, poorly connected and highly nested networks tend to have low coevolutionary temperature. Including the average interaction degree symmetry in the SEM reveals that it mediates the relationship between nestedness and reciprocity: nestedness is negatively associated with the average interaction degree symmetry which, in turn, is positively associated with reciprocity (Figure S3). We also find significant positive, although weaker, relationship between local network modularity and coevolutionary strength. However, there is no significant association between patch size and modularity.

**Figure 3:**
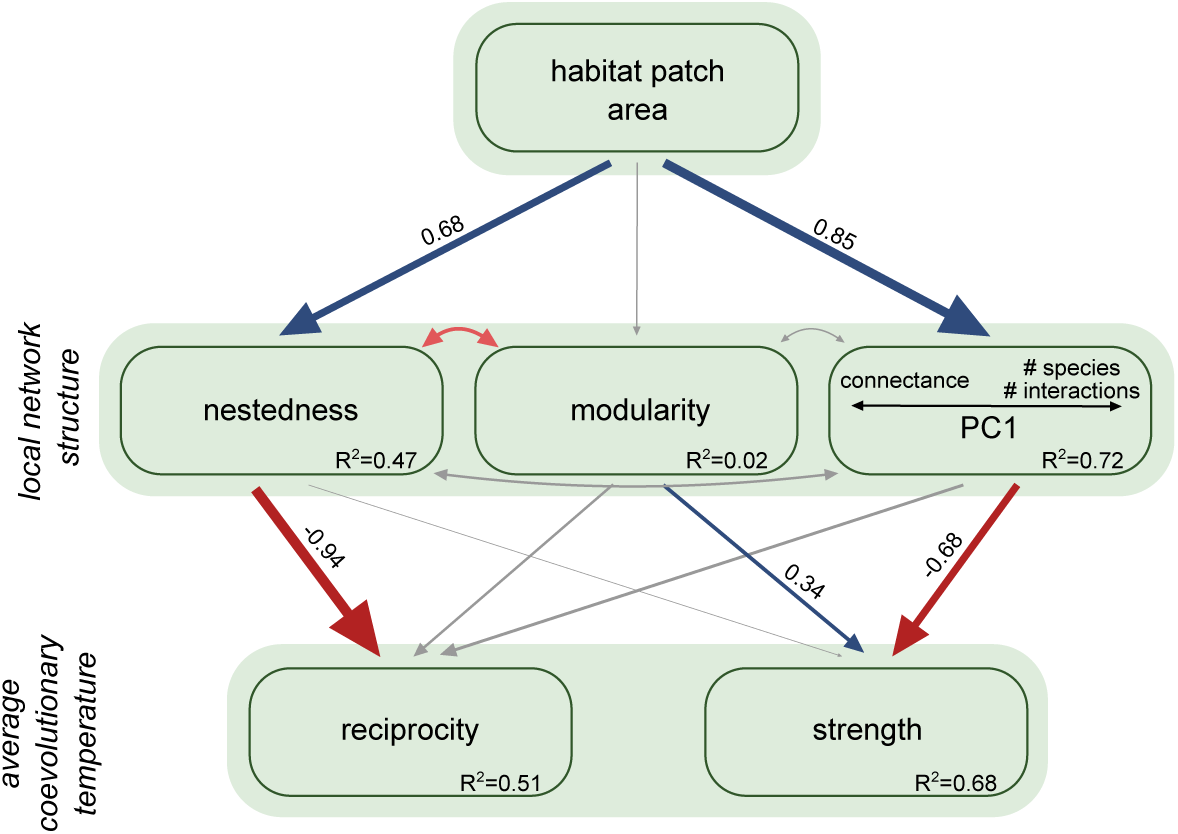
Effect of patch area on local network structure, and coevolutionary reciprocity and strength, as predicted by the structural equation model (SEM). Significant positive and negative relationships (*p <* 0.05) are shown with blue and red arrows, respectively. Nonsignificant relationships are shown with grey arrows. Arrow thickness is proportional to the strength of the relationship. Numbers adjacent to arrows are the standardised path coefficients from the SEM. Double-headed arrows between the network structure measures indicate covariances. Habitat patch area was *ln*-transformed. Nestedness and modularity were standardised as Z-scores using a null model.

At the interaction scale, we find a strong correlation between interaction degree symmetry and reciprocity (Figure 4A). Interactions between species with similar degrees have higher reciprocity, but also higher variability in reciprocity across replicate simulations, than interactions between species with dissimilar degrees. There is no clear relationship between coevolutionary strength, or its variability across replicas, and interaction degree symmetry (Figure 4B). An interaction between the same two species, for example, *K. arvensis* and *E. balteatus*, can vary substantially in interaction degree symmetry, reciprocity, and strength across local networks.

**Figure 4:**
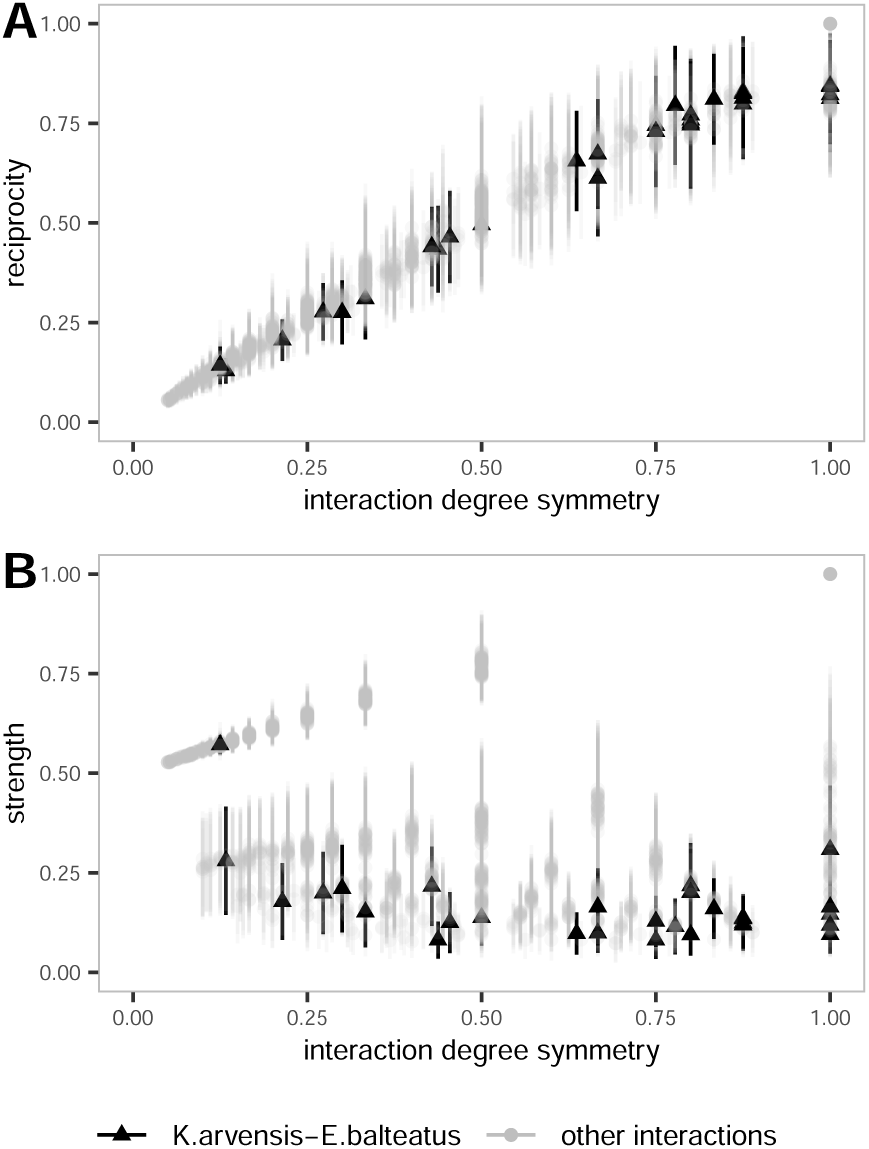
Relationship between interaction degree symmetry and coevolutionary (A) reciprocity and (B) strength. The interaction between *K. arvensis* and *E. balteatus* is indicated by black triangles, whereas all other interactions are shown as grey circles. Points correspond to mean values across replicate simulations for each interaction in each grassland fragment, whereas vertical lines show the standard deviations.

## Discussion

We introduce an approach for assessing coevolutionary temperature in terms of the reciprocity and strength of pairwise coevolutionary effects. We show that coevolutionary temperature of plant–pollinator interactions in a fragmented landscape exhibits a geographic mosaic shaped by the spatial variation in community structure. By investigating the drivers of this variability at both the community and interaction scales, our results provide insights into where and why coevolutionary hotspots and coldspots may arise.

At the community scale, we find that smaller habitat patches tend to harbour communities with higher average reciprocity and average strength, suggesting that such patches may act as coevolutionary hotspots. These communities tend to be small, highly connected and poorly nested, and contain many interactions with high degree symmetry (interactions between species with a similar number of partners, i.e, generalist–generalist or specialist–specialist). We find that high reciprocity is associated with high degree symmetry, which in turn, is driven by low nestedness. Strong coevolutionary effects are associated with small and highly connected networks. Conversely, larger patches support species-rich, nested, and weakly connected communities, resulting in lower reciprocity and strength, and thus, are more likely to act as coldspots.

At the interaction scale, we find a strong positive relationship between degree symmetry and reciprocity. This result highlights how the role of species within the network can shape the likelihood of their interactions functioning as hotspots. Importantly, the same pair of interacting species can exhibit different coevolutionary temperatures across fragments, depending on the local community. This context dependence echoes the predictions of the GMTC by emphasizing that hotspots and coldspots are not intrinsic properties of particular species pairs, but instead emerge from the interplay of community composition and landscape setting (Thompson et al., 2017; Thompson, 2005).

Understanding where in the landscape, and for which interactions, hotspots and coldspots are most likely to emerge has important implications for conservation and restoration planning. Changes in species composition, whether through extinctions or introductions, can reconfigure a network, and thus, alter the coevolutionary temperature. Our results suggest that habitat loss and fragmentation may increase the likelihood of hotspots, and thus drive divergence in coevolutionary trajectories among populations (Costa et al., 2025; Gawecka et al., 2022; Faillace et al., 2021). Such differences could, in turn, complicate the recovery of communities after restoration (Evans et al., 2022; LaRue et al., 2017). Depending on goals, restoration strategies might either foster hotspots by creating small patches, or promote coldspots by expanding existing habitats.

The role of habitat area in shaping coevolution has been examined empirically by Siepielski and Benkman (2005) in a pairwise antagonistic system between crossbills and lodgepole pines. In contrast to our findings, they reported that larger habitat patches acted as hotspots: crossbill densities increased with forest area, intensifying reciprocal selection on pine seed defences and crossbill bill morphology. This discrepancy highlights two important considerations. First, habitat area may influence coevolution in mutualistic and antagonistic systems in different ways. For example, habitat area has been shown to affect primarily modularity of antagonistic networks (Grass et al., 2018; Hagen et al., 2012). Yet, in our mutualistic context, modularity is associated with coevolutionary strength, but not reciprocity (Figure 3). Second, the pattern in Siepielski and Benkman (2005) suggests that species abundance, and thus interaction frequency, is a key determinant of selection strength. Yet, our model treats interactions as unweighted. Incorporating interaction frequencies through weighted networks is a promising direction for extending our framework.

Here, we focus on patch size as a driver of community structure, and consequently, coevolutionary effects. However, other landscape characteristics such as habitat quality, matrix composition, patch connectivity, or spatial configuration are known to influence community composition and structure (e.g., Bradfer-Lawrence et al., 2025; Waddell et al., 2024; Ryser et al., 2019). For example, Gilarranz et al. (2015) showed that peripheral patches harbour smaller and less nested mutualistic networks than central patches. Ren et al. (2023) found that plant–pollinator networks in forest interiors are smaller and less nested than those at forest edges. When viewed alongside our results, these patterns raise the possibility that peripheral patches and forest interiors are more likely to act as coevolutionary hotspots.

Our analyses rely on static community snapshots derived from observed plant–pollinator interactions. Consequently, we do not capture eco-evolutionary processes involving dispersal, colonization–extinction dynamics, or ongoing trait remixing (Legrand et al., 2017; Toju et al., 2017; Urban et al., 2008). Yet, these processes may cause coevolutionary temperature to fluctuate not only across space but also through time (Thompson, 2005). Applying our framework to eco-evolutionary models (Gawecka et al., 2022; Govaert et al., 2019; Andreazzi et al., 2018) could provide insights into the temporal dependence of hotspots and coldspots.

Finally, we emphasise that our proposed measures of reciprocity and strength derive from a specific coevolutionary model (Guimarães et al., 2017), and thus, represent one possible way to quantify coevolutionary effects. Other theoretical or empirical approaches may define and measure coevolution differently, which could lead to alternative patterns or interpretations. Importantly, our measures have not yet been validated empirically. Because quantifying coevolution in the field is challenging, especially in species-rich communities, most empirical studies rely on measuring trait convergence or divergence. However, such trait-based signals do not always correspond directly to the magnitude or reciprocity of co-evolutionary effects, particularly in a community setting. Indirect approaches that combine mechanistic models with statistical methods (Nuismer et al., 2022; Week and Nuismer, 2019) offer a promising path forward for bridging these perspectives and, ultimately, evaluating the generality of our findings.

We highlight that advancing our understanding of the geographic mosaic of coevolution in complex systems will require shifting from binary hotspot–coldspot classifications to a continuous gradient of coevolutionary temperature, and from pairwise interactions to species-rich communities. While our study provides a first step toward understanding the mechanisms that generate hotspots and coldspots, we hope it will stimulate further discussion and inspire future research.

## Acknowledgements

We thank Frank Jauker and Ingo Grass for sharing the empirical data, as well as John Thompson and Leandro Cosmo for discussions.

## Data availability

All code used in this study is available on GitHub (github.com/kgawecka/coevolutionary_ temperature).

