## Supporting Information for "Quantifying the geographic mosaic of coevolutionary temperature: from coldspots to hotspots"

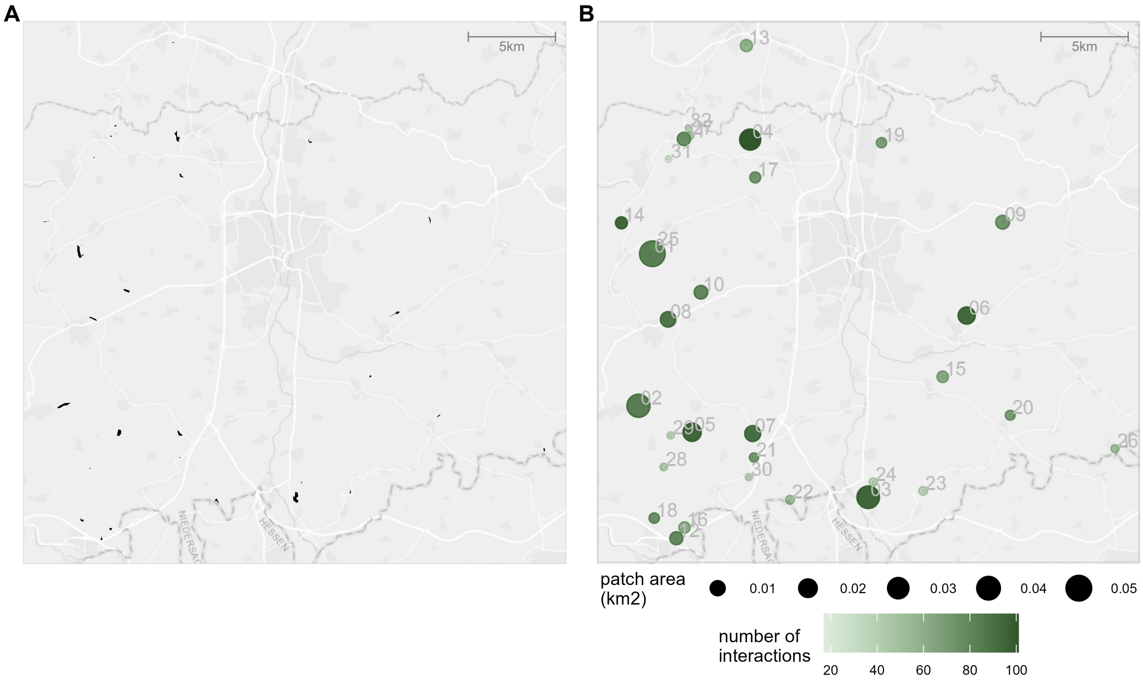


Figure S1: Map of 32 calcareous grassland fragments. (A) Actual locations, shapes and sizes of the grassland fragments. (B) Plot illustrating patch sizes (point size) and the number of plant-pollinator interactions observed (colour).

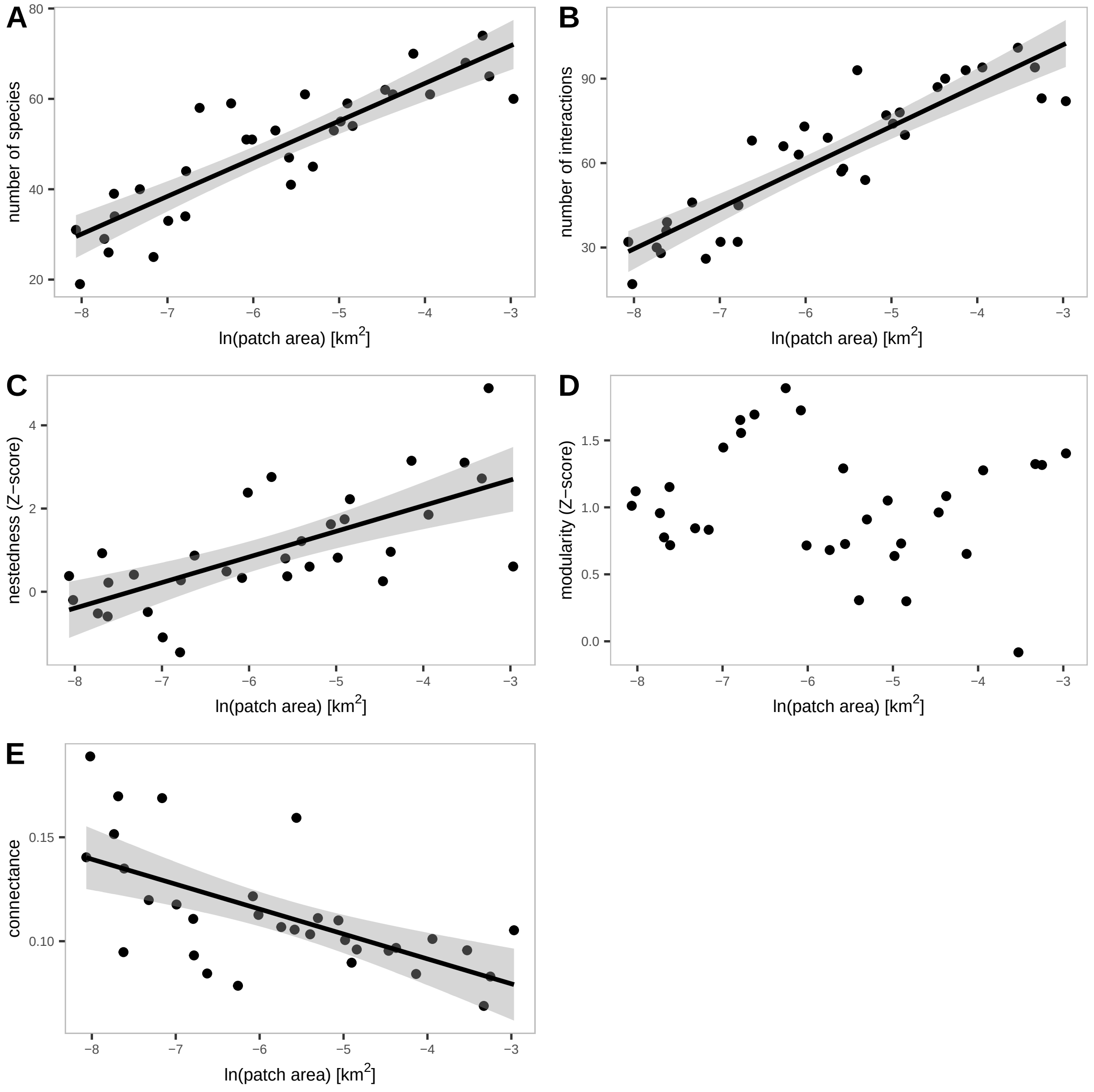


Figure S2: Effect of patch area on local plant-pollinator network structure: (A) number of species, (B) number of interactions, (C) nestedness (as standardised Z-score), (D) modularity (as standardised Z-score), (E) connectance. Points plot the empirical data. Statistically significant relationships (p-value<0.05) are shown with the linear model fit and its 95% confidence interval.

Table S1: SEM coefficients for the model shown in the main text. Patch area is *ln*-transformed; nestedness and modularity are expressed as Z-scores from probabilistic null models.

| Response | Predictor | Estimate | Std. Error | DF | *P* | Std. Est. |
|---|---|---|---|---|---|---|
| PC1 | patch area | 0.932 | 0.105 | 30 | $< 0.001$ | 0.850 |
| nestedness | patch area | 0.616 | 0.120 | 30 | $< 0.001$ | 0.684 |
| modularity | patch area | -0.044 | 0.053 | 30 | 0.409 | -0.151 |
| R_mean | PC1 | 0.011 | 0.007 | 28 | 0.138 | 0.282 |
| R_mean | nestedness | -0.046 | 0.010 | 28 | $< 0.001$ | -0.939 |
| R_mean | modularity | -0.035 | 0.022 | 28 | 0.115 | -0.238 |
| S_mean | PC1 | -0.027 | 0.006 | 28 | $< 0.001$ | -0.676 |
| S_mean | nestedness | -0.003 | 0.008 | 28 | 0.679 | -0.067 |
| S_mean | modularity | 0.052 | 0.018 | 28 | 0.007 | 0.345 |
| PC1 ~~ nestedness | | 0.233 | | 32 | 0.104 | 0.233 |
| PC1 ~~ modularity | | 0.141 | | 32 | 0.224 | 0.141 |
| nestedness ~~ modularity | | -0.350 | | 32 | 0.027 | -0.350 |

**$R^2$**

PC1 = 0.72; nestedness = 0.47; modularity = 0.02; R_mean = 0.51; S_mean = 0.68

**Global fit indices**

Chi-squared (df = 3) = 6.401; Fisher's C (df = 6) = 8.730

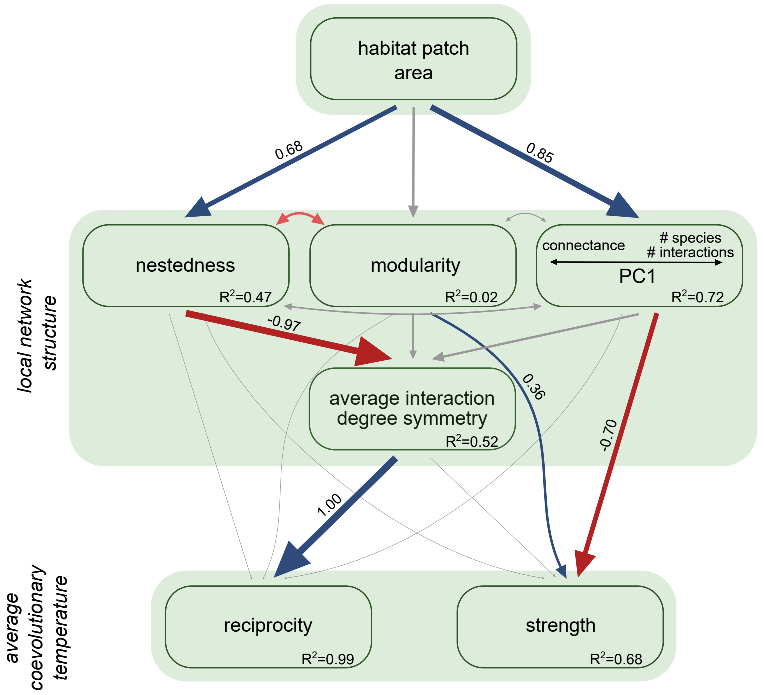


Figure S3: Effect of patch area on local network structure (PC1, nestedness, modularity and average interaction degree symmetry), and coevolutionary reciprocity and strength, as predicted by the SEM. Significant positive and negative relationships ($p < 0.05$) are shown with blue and red arrows, respectively. Nonsignificant relationships are shown with grey arrows. Arrow thickness is proportional to the strength of the relationship. Numbers adjacent to arrows are the standardised path coefficients from the SEM.

Table S2: SEM coefficients for the alternative model (Figure S3). Patch area is *ln*-transformed; nestedness and modularity are expressed as Z-scores from probabilistic null models.

| Response | Predictor | Estimate | Std. Error | DF | *P* | Std. Est. |
|---|---|---|---|---|---|---|
| PC1 | patch area | 0.932 | 0.105 | 30 | $< 0.001$ | 0.850 |
| nestedness | patch area | 0.616 | 0.120 | 30 | $< 0.001$ | 0.684 |
| modularity | patch area | -0.044 | 0.053 | 30 | 0.409 | -0.151 |
| degree symmetry | PC1 | 0.014 | 0.008 | 28 | 0.081 | 0.331 |
| degree symmetry | nestedness | -0.049 | 0.010 | 28 | $< 0.001$ | -0.972 |
| degree symmetry | modularity | -0.037 | 0.023 | 28 | 0.107 | -0.242 |
| reciprocity | degree symmetry | 0.958 | 0.025 | 27 | $< 0.001$ | 1.003 |
| reciprocity | PC1 | -0.002 | 0.001 | 27 | 0.072 | -0.049 |
| reciprocity | nestedness | 0.002 | 0.002 | 27 | 0.343 | 0.035 |
| reciprocity | modularity | 0.001 | 0.003 | 27 | 0.848 | 0.004 |
| strength | degree symmetry | 0.064 | 0.150 | 27 | 0.676 | 0.066 |
| strength | PC1 | -0.028 | 0.006 | 27 | $< 0.001$ | -0.698 |
| strength | nestedness | -0.000 | 0.011 | 27 | 0.992 | -0.002 |
| strength | modularity | 0.054 | 0.019 | 27 | 0.008 | 0.361 |
| PC1 $\sim\sim$ nestedness | | 0.233 | | 32 | 0.104 | 0.233 |
| PC1 $\sim\sim$ modularity | | 0.141 | | 32 | 0.224 | 0.141 |
| nestedness $\sim\sim$ modularity | | -0.350 | | 32 | 0.027 | -0.350 |

**$R^2$**

PC1 = 0.72; nestedness = 0.47; modularity = 0.02; degree symmetry = 0.52; reciprocity = 0.99; strength = 0.68

**Global fit indices**

Chi-squared (df = 4) = 9.503; Fisher's C (df = 8) = 14.028

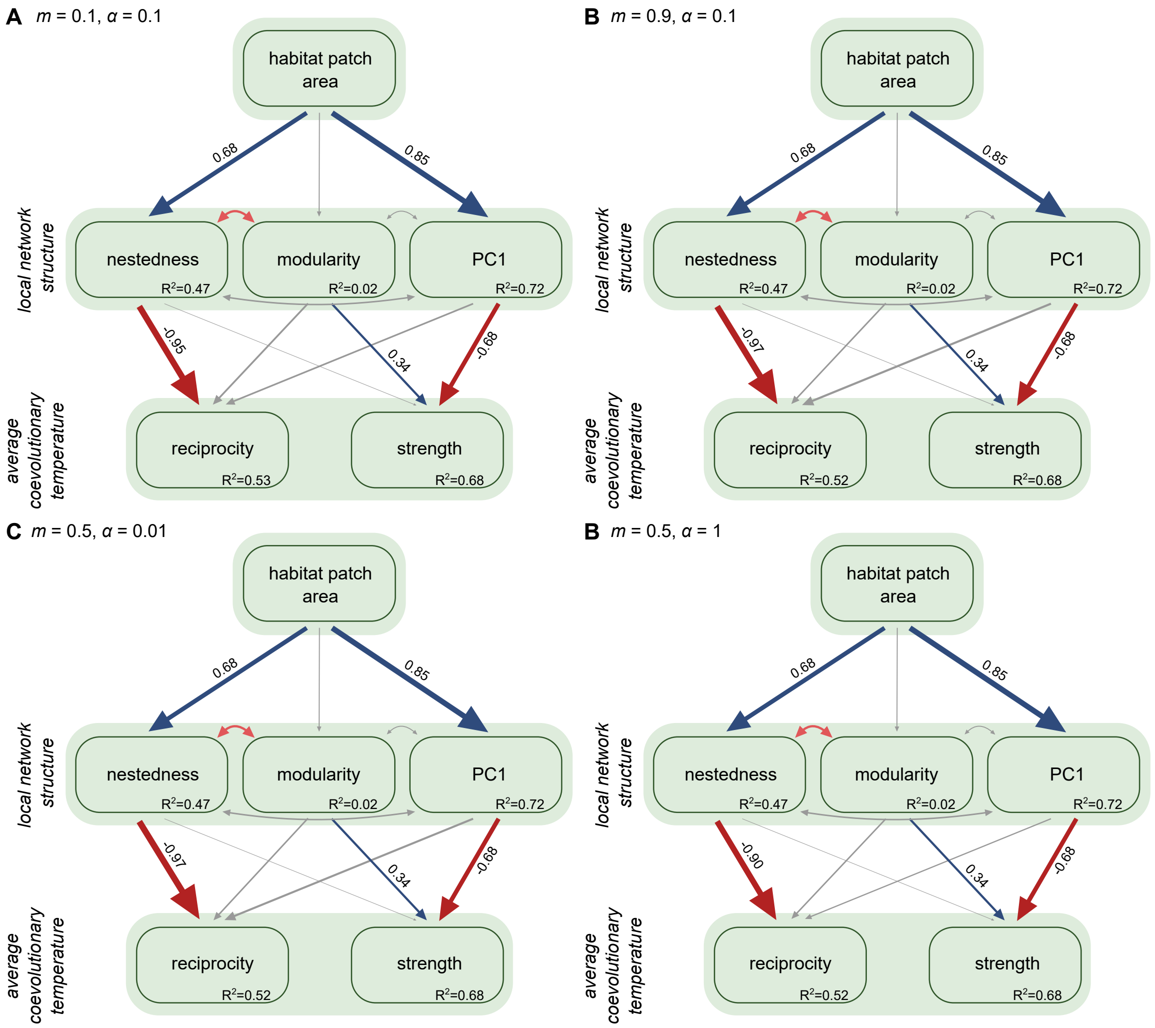


Figure S4: Effect of patch area on local network structure (PC1, nestedness and modularity), and coevolutionary reciprocity and strength, as predicted by the SEM. Significant positive and negative relationships ($p < 0.05$) are shown with blue and red arrows, respectively. Nonsignificant relationships are shown with grey arrows. Arrow thickness is proportional to the strength of the relationship. Numbers adjacent to arrows are the standardised path coefficients from the SEM. (A) $m = 0.1$, $\alpha = 0.1$, (B) $m = 0.9$, $\alpha = 0.1$, (C) $m = 0.5$, $\alpha = 0.01$, (D) $m = 0.5$, $\alpha = 1$.

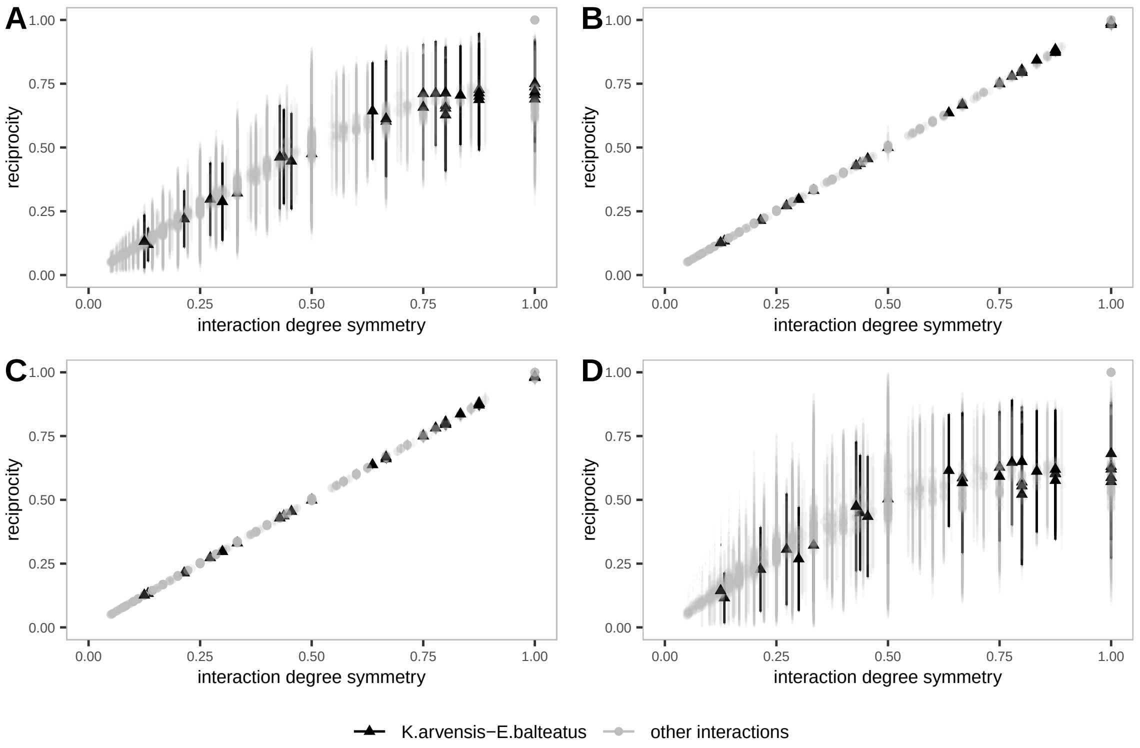


Figure S5: Relationship between interaction degree symmetry and coevolutionary reciprocity. The interaction between *K. arvensis* and *E. balteatus* in indicated by a black triangles, whereas all other interactions are shown as grey circles. Points correspond to mean values across replicate simulations for each interaction in each grassland fragment, whereas vertical lines show the standard deviations. (A) $m = 0.1$, $\alpha = 0.1$, (B) $m = 0.9$, $\alpha = 0.1$, (C) $m = 0.5$, $\alpha = 0.01$, (D) $m = 0.5$, $\alpha = 1$.

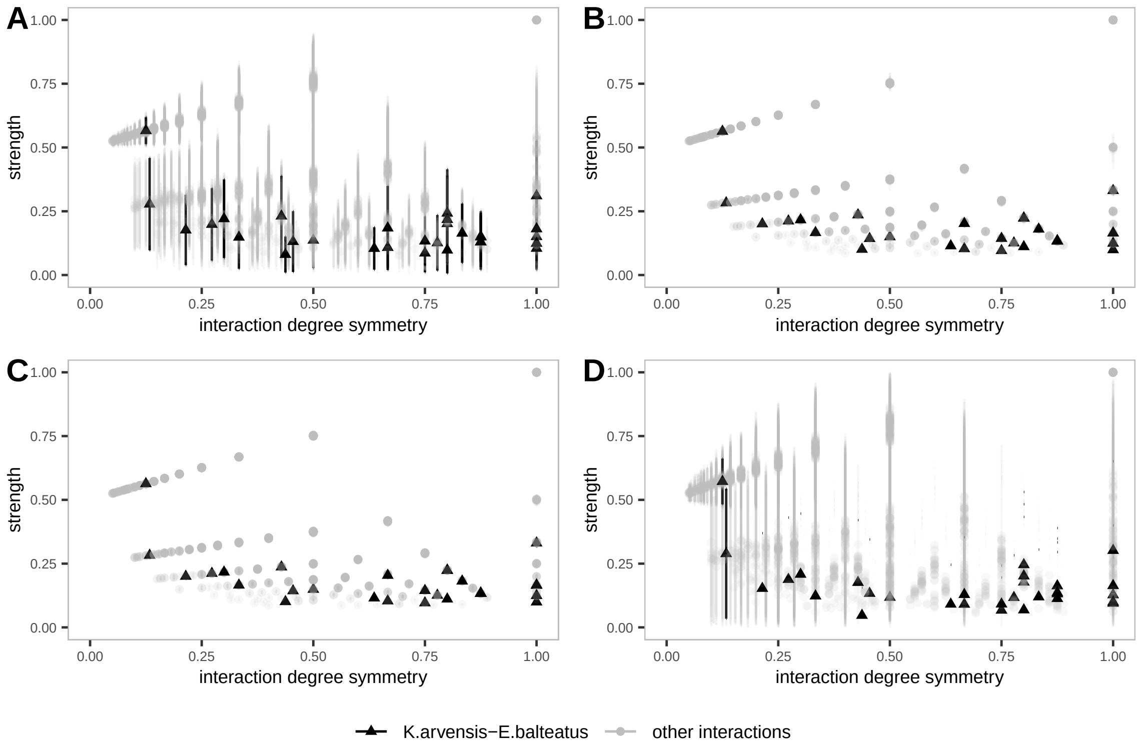


Figure S6: Relationship between interaction degree symmetry and coevolutionary strength. The interaction between *K. arvensis* and *E. balteatus* in indicated by a black triangles, whereas all other interactions are shown as grey circles. Points correspond to mean values across replicate simulations for each interaction in each grassland fragment, whereas vertical lines show the standard deviations. (A) $m = 0.1$, $\alpha = 0.1$, (B) $m = 0.9$, $\alpha = 0.1$, (C) $m = 0.5$, $\alpha = 0.01$, (D) $m = 0.5$, $\alpha = 1$.
